# Three-dimensional dipole orientation mapping with high temporal-spatial resolution using polarization modulation

**DOI:** 10.1101/2023.12.12.571225

**Authors:** Suyi Zhong, Liang Qiao, Xichuan Ge, Xinzhu Xu, Yuzhe Fu, Shu Gao, Karl Zhanghao, Huiwen Hao, Wenyi Wang, Meiqi Li, Peng Xi

## Abstract

Fluorescence polarization microscopy is widely used in biology for molecular orientation properties. However, due to the limited temporal resolution of single-molecule orientation localization microscopy and the limited orientation dimension of polarization modulation techniques, achieving simultaneous high temporal-spatial resolution mapping of the three-dimensional (3D) orientation of fluorescent dipoles remains an outstanding problem. Here, we present a super-resolution 3D orientation mapping (3DOM) microscope that resolves 3D orientation by extracting phase information of the six polarization modulation components in reciprocal space. 3DOM achieves an azimuthal precision of 2° and a polar precision of 3° with spatial resolution of 128 nm in the experiments. We validate that 3DOM not only reveals the heterogeneity of the milk fat globule membrane, but also elucidates the 3D structure of biological filaments, including the 3D spatial conformation of λ-DNA and the structural disorder of actin filaments. Furthermore, 3DOM images the dipole dynamics of microtubules labeled with green fluorescent protein in live U2OS cells, reporting dynamic 3D orientation variations. Given its easy integration into existing wide-field microscopes, we expect the 3DOM microscope to provide a multi-view versatile strategy for investigating molecular structure and dynamics in biological macromolecules across multiple spatial and temporal scales.

## Main

Fluorescent molecules exhibit dipole behavior and their orientation provides insights into the properties of the targeted molecular [1-3]. Numerous fluorescence polarization microscopes (FPM) [4-7] have been developed to measure dipole orientation, which helps to study the motion of molecular motors [8], conformation of DNA [9, 10], molecular arrangement of actin filaments [11-13], and lipid membranes [14-16], etc. To overcome the challenge of conventional FPM limited by optical diffraction, improved super-resolution FPM techniques have been proposed [17-20], such as single-molecule orientation-localization microscopy (SMOLM) [21-23], and polarization modulation [24]. However, the limited temporal resolution and specialized sample preparation of SMOLM make rapid bioimaging a formidable challenge [25, 26]. While polarization modulation techniques such as super-resolution by polarization demodulation (SPoD) [24] and super-resolution dipole orientation mapping (SDOM) [17], can offer fast imaging speeds to capture the biomolecule dynamics, they can only detect the orientation of the dipole projected on a two-dimensional (2D) plane [27-32]. The three-dimensional (3D) orientation mapping plays a crucial role in mitigating the angular retrieval bias caused by projections in 2D orientation retrieval [33, 34]. For instance, the interaction of labeled F-actin with different proteins will result in 3D conformations with different structures. To distinguish the filament organization or the compactness of the filament bonds, it is crucial to obtain the 3D orientation of the target molecule [35]. However, to rapidly image and resolve 3D orientation simultaneously at resolution beyond diffraction limit, with abundant labeling density to biological samples (unlike single-particle/single-molecule tracking), is rather challenging.

To address this, we have developed 3D orientation mapping (3DOM) microscopy, through widefield (WF) polarization modulation and oblique illumination microscopy. 3DOM successfully demonstrates high-precision 3D orientation quantification together with <100 nm spatial resolution. By encoding the additional orientation dimension in reciprocal space, 3DOM further integrates dipole angle mapping with super-resolution structure. The super-resolution image is effectively reconstructed using the fast iterative shrinkage-thresholding algorithm (FISTA) [36], and the dipole orientation is resolved by employing the excited polarization components in reciprocal space. We derive the fundamental precision limits via Cramér-Rao bound (CRB) [37] and Fisher-information to evaluate the orientation precision, providing insights for optimization of experiments. Experimental validations with both simulated and live cell samples confirm precise 3D molecular orientation determination. We showcase DNA conformational sensing, membrane heterogeneity mapping, and cytoskeletal imaging as prime applications. Compared to existing 2D techniques, 3DOM provides more comprehensive dipole angle measurement to mitigate orientation biases.

Overall, 3DOM expands the super-resolution toolkit with fast nanoscale 3D dipole orientation imaging. It shows immediate potential to uncover previously obscured molecular processes through resolving positional and angular biomolecule dynamics. This work helps enable the next leap for fluorescence polarization techniques towards rapid 3D molecular metrology in cells.

## Results

### Principle of operation and orientation mapping

Details of the customized 3DOM system design are provided in the Methods (Fig. 1a, b, modified from PolarSIM system in Airy Technologies Co.). In the polarization modulation, we use a spatial light modulator to generate diffracted light with different oblique angles. Then the outgoing light is modulated into *p*-polarized light by half-wave plates and polarizing beam splitter, and then a custom-made mask selects the +1-order light to pass through. Finally, the lens converges the +1-order light onto the non-central position in the back focal plane of the objective. To ensure the uniqueness of the solution[38], we set six different positions in the BFP (Fig. 1b), and the positions corresponds to the mask. Benefitted from the digital projection of the liquid crystal on silicon spatial light modulator (SLM), our system allows for a raw image acquisition speed of up to 1697 fps and a 3DOM imaging speed of up to 283 fps (6 raw images per reconstruction).

**Fig. 1.**
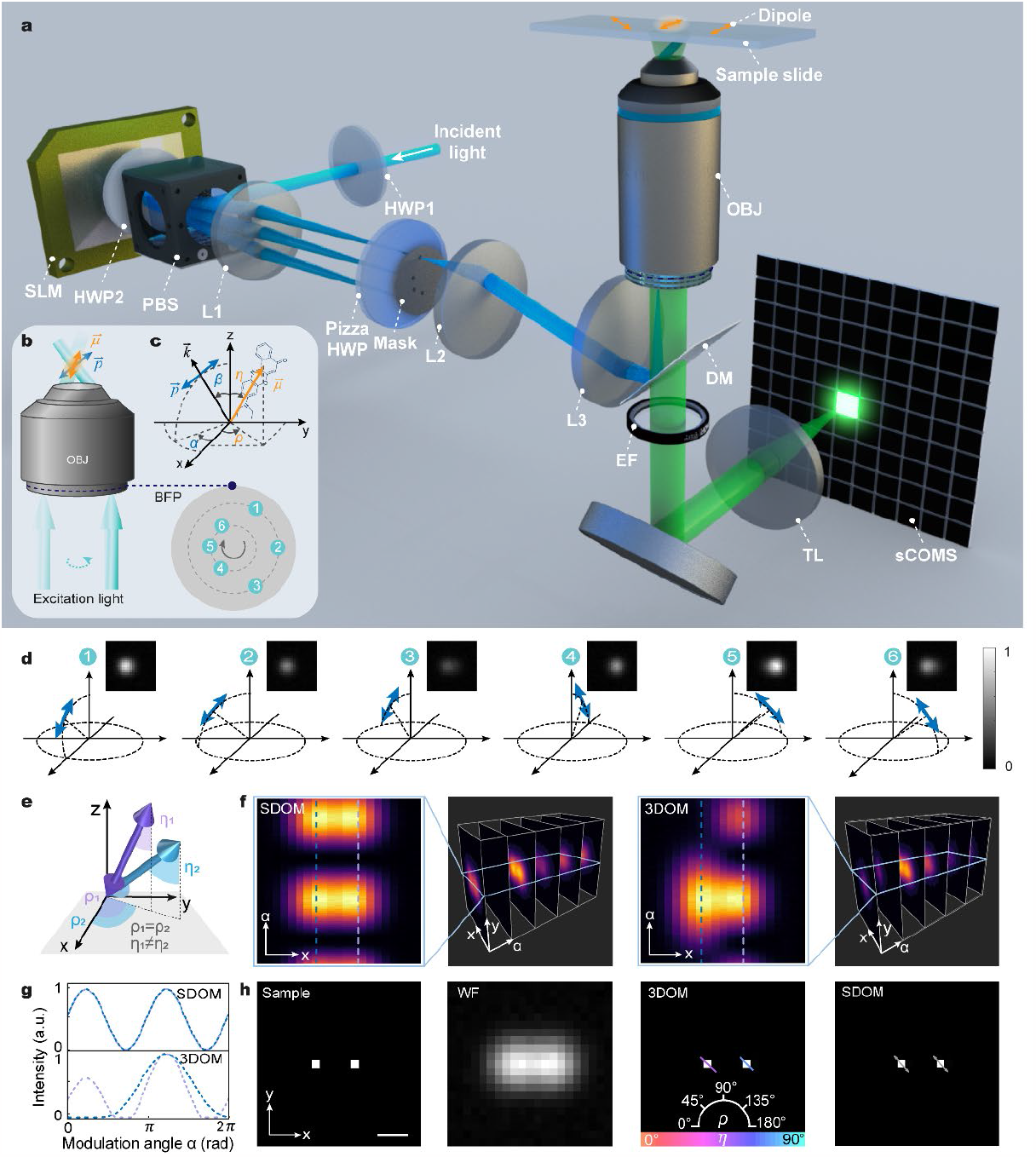
Schematic illustration of the 3D orientation mapping (3DOM) microscopy. **a** Experimental setup of 3DOM. SLM, spatial light modulator; HWP, half-wave plates; PBS, polarizing beam splitter; OBJ, objective; DM, dichroic mirror; EF, emission filter; TL, tube lens. **b** Polarization modulation depicts the excitation beam (green), *p*-polarized light (blue), and dipole emitter 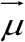 (orange). The excitation light converges at the BFP of the objective lens, and different positions of the convergence change in the excitation light’s outgoing direction and polarization state. **c** Definition of the dipole moment direction 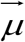 (chemical structure example of a Nile red fluorophore), characterized by its azimuthal angle *ρ* and polar angle *η*. The excitation light is a plane wave with a wave vector 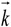, where *p*-polarized light is defined by *α* and *β*. Here, we consider *β* ≠ 0°. **d** The emission intensity distribution under polarized excitation modulation. **e** Schematic diagram of dipoles with the same azimuthal angle, different polar angles. **f, g** Comparison of the emission intensity variations in SDOM and 3DOM for, and a distance of 250 nm. **h** 3DOM resolves both azimuthal angle *ρ* (indicated by the direction of the rod) and polar angle *η* (indicated by the color), with length proportional to sin *η*. In contrast, SDOM only yields an azimuthal angle (indicated by the gray arrow). Scale bar = 200 nm.

In Fig. 1c, we represent the fluorescent molecule by the electric dipole moment along the unit vector in spherical coordinates, which is expressed as 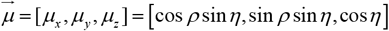, where *ρ* and *η* are the azimuthal angle and polar angle of 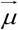. In this work, azimuthal angle refers to the clockwise angle between the projection of the vector on the *x-y* plane and the *x*-axis, and polar angle is the angle between the vector and the *z*-axis. The polarization state of the excitation light is characterized as 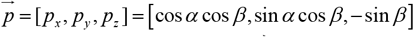, where *α* and *β* to define the azimuthal angle and oblique angle of 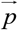, respectively.

The effective photons emitted by each fluorescent dipole are quantified by the cosine square of the angle [39], denoted by 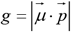. Six-fold polarization-modulated illumination induces observable changes in the polarization dimension in the reciprocal space. Then, we expand the pixel level intensity expression obtained for the *m*th polarization modulation as follows:

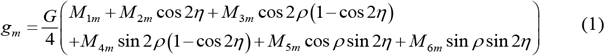

where *G* represents the sample irradiance, which is related to exposure time and molecular quantum efficiency. *M*_1*m*_ ∼ *M*_6*m*_ are describing the polarization state of the excited light. The fluorescence image *I*_1_ ∼ *I*_6_ captured images under six different polarization modulations can be represented as a set of polarization response equations (Fig. 1d). By applying the inversion of the coefficient *M* = [*M*_*im*_]_6×6_, we can combine the dipole orientation information in the reciprocal space (for details, see Supplementary Section 1), resulting in the following expression:

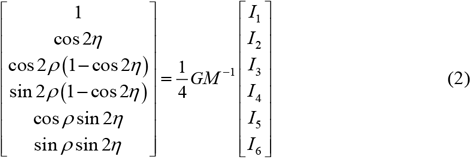

By changing the polarization state of the excitation light six times, 3DOM can distinguish the different dipoles with 3D orientation in the polarization domain, even for dipoles with different angles to the *z*-axis (Fig. 1e-g, see Supplementary Section 2 for comparison of different orientations of the dipoles), while SDOM only yields 2D orientation (Fig. 1h and Supplementary Fig. S1).

### Theoretical performance on precision and angular accuracy

To assess the optimal estimation precision of the orientation in the 3DOM model, we derive the Fisher-information and the CRB for quantitative evaluation (for details, see Supplementary Section 3). Assume that the orientation of a single fluorescent dipole is Ω(*ρ, η*), the normalized fluorescence intensity collected at the *m*th polarization modulation is *I*_*m*_ (Ω), and the number of photons reaching the *n*th pixel is *N*_*mn*_ (Ω). We mainly consider Poisson noise and Gaussian noise affecting the output signal. The Poisson distribution *P*_*s*_ (*N*_*mn*_) characterizes the statistical uncertainty of photons reaching the detector and the Gaussian distribution *P*_*g*_ (*N*_*mn*_) depicts the readout noise associated with the detector. Therefore, the detector readout value *q*_*mn*_ follows the probability distribution:

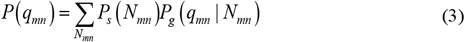

Then, the total Fisher-information *F*_ΩΩ_ can be obtained by summing the individual pixel Fisher-information *F*_*mn*_ as follows:

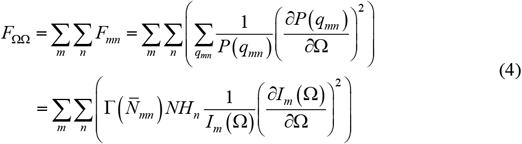

Here, we define a weighting factor 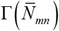 for Gaussian and Poisson noise, with a factor value closer to 1 indicating less influence of noise. In the absence of additional noise, combining the precision of the azimuthal angle and the polar angle gives the solid angle orientation precision [38], the CRB for unbiased estimation can be obtained by:

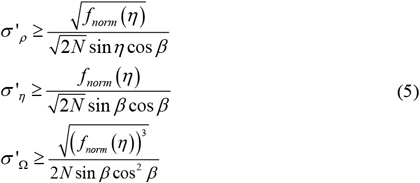

Noise is unavoidable in measurements. We can compute the Fisher-information considering Poisson and Gaussian noise as follows:

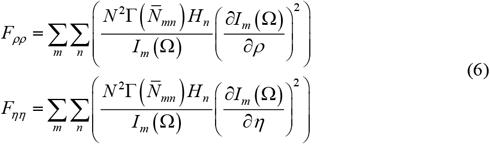

Subsequently, the CRB for unbiased estimation, considering the additional noise, serves as the bound for orientation precision, as follows:

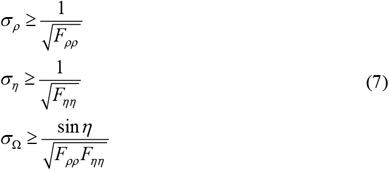

The precision limits of azimuth and polar angle are dependent on the polar angle of dipole, noise level, total number of photons, and incidence angle of excitation light. Fig. 2a illustrates the CRB for the azimuthal, polar, and solid angles at various oblique incidence angles *β* and dipole polar angles *η*. The calculations consider a total number of photons *N* of 2000 and a noise level *σ*_g_ of 50. The fluctuation of the light intensity changes as *β* changes (Supplementary Fig. S2). When *β* is less than 45°, increasing dipole *η* from 45° to 75° boots intensity but reduces sensitivity, resulting in lower precision. Conversely, decreasing the *η* between 15°and 45° yields higher precision, despite the lower intensity. The results are opposite when the *β* is larger than 45° (Fig. 2a). Additionally, the precision *σ*_*ρ*_ sin *η* worsens with a decrease in *N* and an increase in *σ*_g_ (Fig. 2d). This effect is particularly significant for dipoles with high polar angles. The precisions *σ*_*η*_ and *σ*_Ω_ worsen with lower *N* and higher *σ*_g_, especially for dipoles with high *η* between 30° and 70°. To maintain *σ*_Ω_ within 5 sr, the ratio of *N* to *σ*_g_ should be greater than 25. For instance, with *N* at 2000 and *σ*_g_ at 50, the precisions for 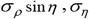 and *σ*_Ω_ are 1.51°, 1.08° and 1.63 sr, respectively.

**Fig. 2.**
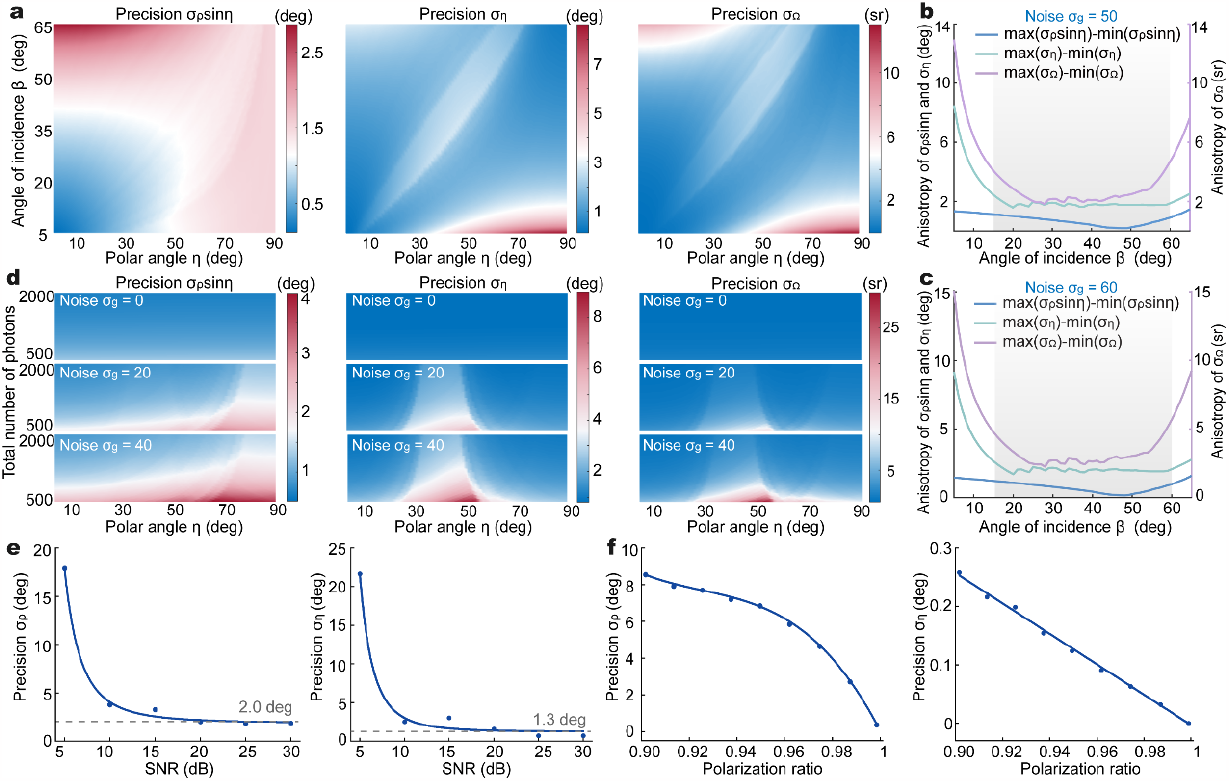
The influence of different factors on polarization resolving precision. **a** The effect of the excitation oblique angle *β* on the Cramér-Rao bound (CRB). The azimuth precision is scaled by the sin *η* factor, with a total photon number *N* of 2000 and noise *σ*_*g*_ of 50. **b** The anisotropy of the dipole precision varies with the oblique angle *β* in (**a**). The anisotropies of precisions are calculated by subtracting the maximum precision value from the minimum precision value at a specific angle. The gray area is the recommended range for *β* = 15°∼60°. **c** The variation of anisotropy of *σ*_*ρ*_ sin *η, σ*_*η*_ and *σ*_Ω_ with the oblique angle *β* under a total photon number *N* of 2000 and noise *σ*_*g*_ of 60. **d** Variations of orientation precisions with the total number of photons and polar angle. All plots are computed with *β* = 35° and noise *σ*_*g*_ = 0, 20 and 40, respectively. **e** The variation curves of the precision *σ*_*ρ*_ and *σ*_*η*_, respectively, with different SNRs of raw data. **f** The impact of polarization ratio on the precision *σ*_*ρ*_ and *σ*_*η*_, respectively. The full width at half maximum (FWHM) of the PSF is set to 250 nm.

We compare the anisotropy in precision for different oblique incidence angles under *N* of 2000 and different noises. It is possible to achieve nearly isotropic precision even without prior knowledge of the dipole orientation distribution within the range of *β* from 15° to 60°, with the anisotropy of 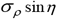 and *σ*_*η*_ both can be within ∼5° and the variation in *σ*_Ω_ is ∼2 sr (Fig. 2b, c). In the absence of noise, isotropic precision can be achieved with set to 35° (Supplementary Fig. S3).

To further quantify the effect of the orientation solving process on the orientation precision, we conduct simulation with different SNR and polarization ratio. The 1000 sample points are randomized orientation (*ρ* ∈[0,*π*], *η* ∈[0,*π/*2]) and added with Gaussian and Poisson noise (Fig. 2e, f). When SNR in the raw image increases, orientation precision improves significantly, with reliable estimation possible at SNR>10 dB, and the precision *σ*_*ρ*_ can reach 1.97° and *σ*_*η*_ of 1.25° at SNR>20 dB. As for polarization ratio, we calculate from the polarization extinction ratio *κ* of the device, denoted by (*κ* −1)/(*κ* +1). The precision increases as the polarization ratio rises from 0.90 to 1, particularly when the polarization ratio surpasses 0.95, signifying a substantial precision boost. Therefore, careful adjustment of the linear polarization of incident light, especially considering its impact on *σ*_*ρ*_. In 3DOM, the extinction ratio can be relaxed to 39:1 for excitation light, and depolarization effects on precision can be disregarded when polarization ratio exceeds 0.95.

To evaluate the orientation mapping accuracy of reconstructed 3DOM images, we simulate biological samples with diverse morphologies in Fig. 3. We first simulate a hemisphere similar to the one wrapped with lipid membranes [40], setting the probe molecules perpendicular to the sphere surface and pointing to the sphere center (Fig. 3a and Supplementary Video S1) [41]. In each z-layer, we obtained the *ρ* results as shown in Fig. 3c, and the color varied with the *η*. The layer-by-layer statistics show that the solved orientation values agree with the preset values, with deviations ranging from 0.17° to 7.15° (Fig. 3b), demonstrating high accuracy.

**Fig. 3.**
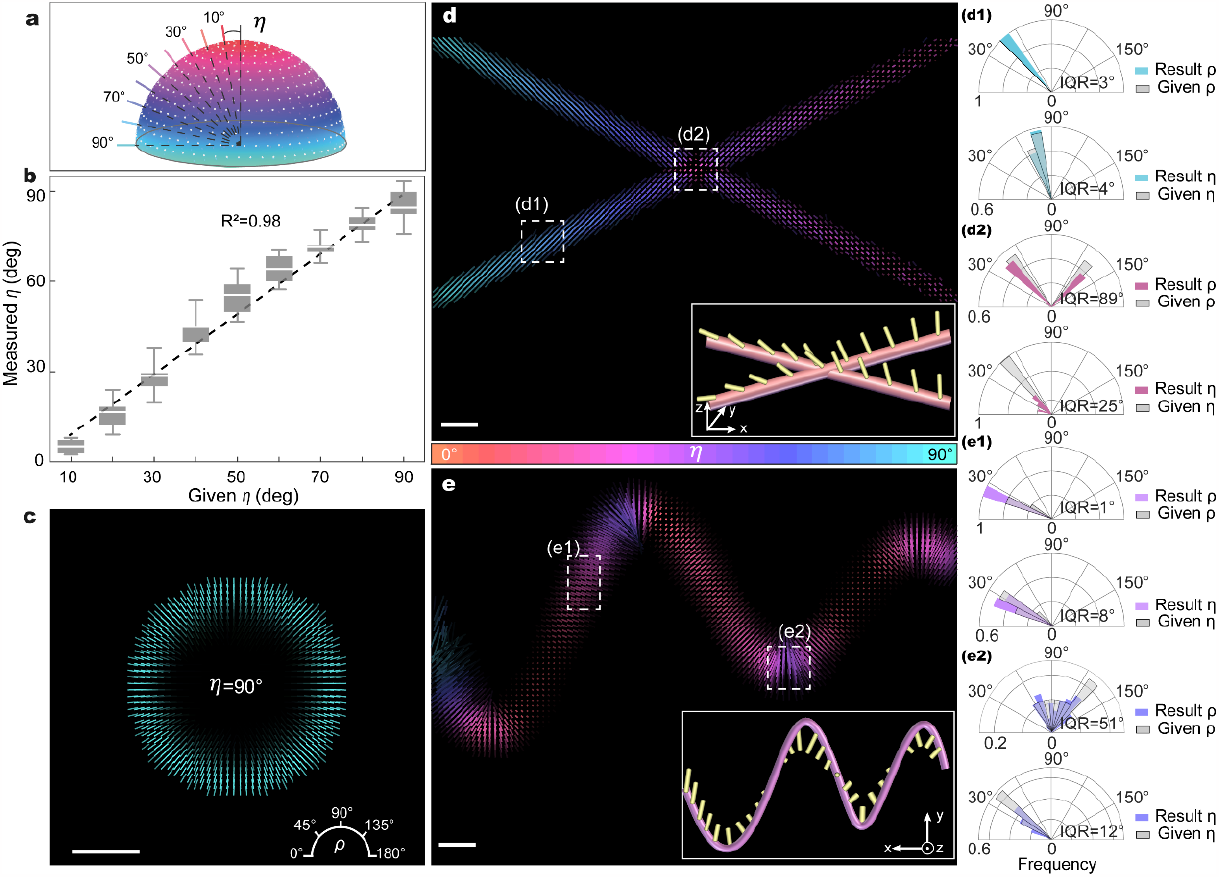
Orientation analysis of simulated biological samples in 3DOM. **a** The schematic illustration of the simulated hemispherical samples. **b** The difference between the 3DOM reconstruction results and the preset angles for different layers of hemispherical samples. The gray dashed line represents the preset value, and the box whisker length and white line represent the standard deviation and mean value, respectively. **c** 3DOM imaging of monolayer sample (polar angle *η* of 90°). **d** 3DOM imaging of the simulated curve sample. The inset shows the distribution settings of the dipoles on the curve. **d1** and **d2** are the statistics distribution of the azimuthal and polar angles, respectively, within the white dashed box in (**d**). **e** 3DOM imaging of the simulated cross-line sample. The inset shows the distribution of the dipoles on the cross-line. **e1** and **e2** are the statistics of the azimuthal and polar angles, respectively, within the white dashed box in (**e**). The interquartile range (IQR) of the reconstruction data is displayed below the respective polar histograms. Rods are orientated according to the solved results *ρ* and color-coded according to the solved results *η*, with their lengths proportional to sin*η*. The FWHM of the PSF is set to 250 nm. *β* set of 35°, the mean value of Gaussian noise to be 0.15% of the fluorescence intensity, and the per-pixel fluorescence intensity as a parameter for the Poisson distribution. Scale bar: 500 nm.

Then, we simulated line samples to understand fibrous structures (insets of Fig. 3d, e). In the simulated curves, the probe molecules are vertically entangled, and in the straight lines, the probe molecules are embedded at different inclination angles (10°∼90°) and the projections are parallel to the straight lines. We quantitatively compare the reconstructed with the preset inter-quadratic range (IQR) of the target region using polar histograms (see Supplementary Table S1 in Supplementary Section 4). The reconstructed 3DOM orientation mappings are shown in Fig. 3d, e, revealing that angular deviations are within ∼4° for straight lines, but curved structures lead to greater deviations with minimal polar angle changes. When lines are closer than 130 nm, orientation mapping bias increases due to light intensity superposition, especially *η*.

We theoretically assess the resolution of adjacent dipoles at different distances and orientations in Fig. 4a-d. When both Δ*ρ* and Δ*η* reach 90°, the deviation in *ρ* and *η* is reduced to within 5°. Additionally, the angular detection deviation decreases as Δ*ρ* increases, which can be explained by the weakening of the modulation capacity on each dipole by the excitation light, leading to a lower image sparsity, ultimately diminishing the accuracy of the reconstruction results. Additionally, a comprehensive depiction of the influence of angular differences on resolution resolves that the resolvable distance improves with both Δ*ρ* and Δ*η* increase.

**Fig. 4.**
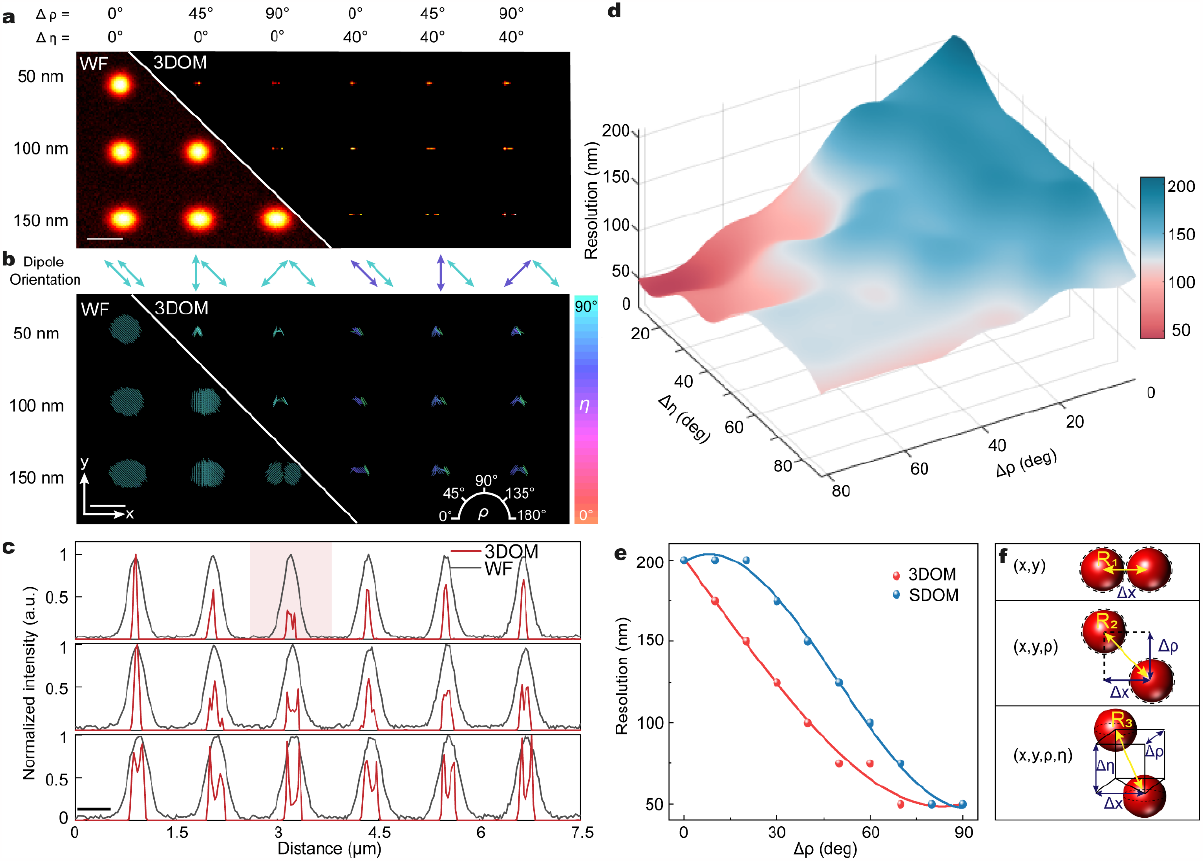
Resolution evaluation in 3DOM. **a** Compare the light intensities of WF and 3DOM at different distances and orientations. The two sample points are convolved with PSF with subjected to background noise, with a SNR of 30 dB. **b** Comparison of the orientation mapping of dipoles. The preset orientation is indicated by the arrows, where the orientation of the arrow represents the azimuthal values and the color represents the polar angle value. **c** The fluorescence intensity profile along the *x*-axis of (**a**), with Δ*ρ* of 90° and Δ*η* of 0° recommended by the red area achieves a resolution of 50 nm. **d** The resolution varies with Δ*ρ* and Δ*η*. **e** A comparison of the resolution between 3D-OM and SDOM at Δ*η* of 0°. **f** Schematic representation of the point distributions of two adjacent points in (*x, y*), (*x, y, ρ*), and (*x, y, ρ, η*) coordinates, respectively. The FWHM of the PSF is set to 250 nm in (**a-e**), and the pixel size is set to 25 nm in (**a-e**). scale bar: 500 nm.

Fig. 4e indicates that both 3DOM and SDOM achieve the highest resolution of 50 nm at Δ*ρ* of 90°, while the resolution of 3DOM surpasses that of SDOM at other angles, which suggests that 3D-OM has stronger resolution in (*x, y, ρ, η*) dimensions. In the absence of polarization considerations (Fig. 4f), the resolution in (*x, y*) coordinate is defined as the minimum distance *R*_1_ at which adjacent points can be distinguished. Considering the azimuthal difference of the dipole, the resolution in (*x, y, ρ*) coordinate is defined by the distance *R*_2_, which includes both the spatial distance and the spacing along the *ρ* -axis. Moreover, when considering the difference in polar and azimuthal angles of the dipoles, the resolution in (*x, y, ρ, η*) coordinate becomes the distance *R*_3_, which includes Δ*x*, Δ*ρ*, and Δ*η*. As the value of Δ*ρ* increases and Δ*x* remains constant, the actual resolution in the (*x, y, ρ*) coordinate becomes higher than that in the (*x, y*) coordinate. Similarly, when Δ*η* increases and keeping Δ*x* and Δ*ρ* constant, the actual resolution in the (*x, y, ρ, η*) coordinate becomes higher than in both the (*x, y, ρ*) and (*x, y*) coordinates. Finally, we demonstrate 3DOM can enhance discrimination capabilities in different dimensions, showing how specific parameter increases result in improved resolution in particular coordinates.

### 3DOM imaging of milk fat globule

Milk fat globules (MFG) have recently attracted greater attention with the growing interest in nutritional, physiological and health properties [42]. Milk fat globule membranes (MFGM) are spatially heterogeneous, thus measurements of MFGM organization and dynamics require specific tool for the study of the 3D orientation [43]. We prepare MFG and label phospholipids using a fluorescent cholesterol analogue with hydrophilic properties, KK114 (for details, see Methods section).

MFG exists in milk in the form of spherical droplets with polydisperse size. Fig. 5 displays a large-sized MFG with a radius of 4.25 μm, potentially resulting from the agglomeration of multiple small MFGs. The MFG core consists mainly of triacylglycerol (TAG), shown as a non-fluorescent black region in the image. We compare the imaging results of 3DOM and pSIM for the organization of phospholipids in MFGM, yielding consistent azimuthal angles. Fig. 5a shows the fluorescence of KK114 in a ring form at the periphery of the MFG, which is characterized along the equatorial section corresponding to the phospholipid markers in the MFGM. In Fig. 5b, the visualization shows the 3DOM results in *x-y, x-z*, and *y-z* views. The colors of the rods in the *x-z* and *y-z* views demonstrate the angle between the dye molecules and the *z*-axis, which also corresponds to the angle between the *z*-axis.

**Fig. 5.**
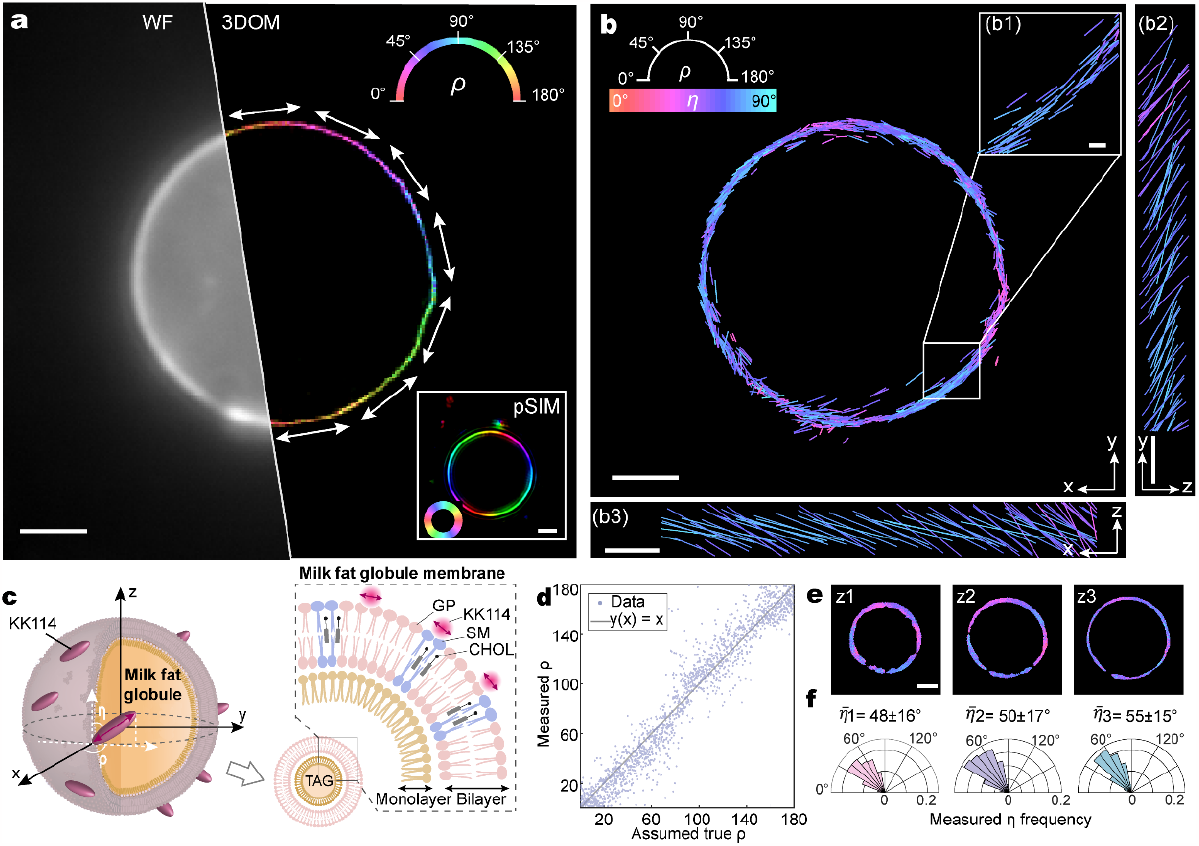
3DOM imaging of milk fat globule (MFG). **a** 3DOM (*ρ* component), pSIM, and WF imaging results of MFG. The white arrows indicate the solved azimuthal orientation along the surface tangent direction of the milk fat globule membrane (MFGM). **b** Visualization of 3DOM imaging results for MFG, with rods orientated and color-coded to indicate *ρ* and*η*, respectively. Zoom in to show the dashed box region and project it in the *x-z* and *y-z* views. **c** Schematic representation of the structure of MFG, which contains a triacylglycerol (TAG) core and a triple-layer structure. KK114 dye labels the phospholipid on the MFGM. GP, glycerophospholipids; SM, sphingomyelin; CHOL, cholesterol. **d** The experimental deviation curve for *ρ* in **(b)** is estimated based on the MFGM tangent direction. **e, f** Imaging and the statistics distribution of *η* orientation within *z*-slice with spacing steps at 1 μm. Scale bars **(a, b, e)**, 2 μm; **(b1-b3)**, 200 nm.

Based on this dataset of Fig. 5b, we generate experimental accuracy curves for azimuthal angle *ρ* (Fig. 5d). By visualizing the orientation of the dye molecules from *x-z* and *y-z* views and performing a 3D observation of the MFG by recording light slices at different *z* depths (Fig. 5e). We histogram the distribution of the polar angle *η* of the dye molecules in the three different layers, and the average *η* orientation of the dye molecules concerning the *z*-axis are 48°±16°, 50°±17° and 55°±15° (median ± s.d.), respectively. We attribute this finding to the fact that the fluorophore is kept out of the hydrophobic bilayer. However, the median absolute deviation of the measurements is larger than the precision of our measurement method. We estimate that this “broadening” of the orientation distribution may be due to the presence of fluorophore linkers, which increase the flexibility of the probe. Next, we speculate a model for probe labeling of MFG based on the statistical orientation of the molecule as shown in Fig. 5c. The resulting MFG products with different particle sizes or different compositional contents present different physicochemical properties. Furthermore, KK114 exhibits liquid-ordered (Lo) preference in both giant unilamellar vesicles and giant plasma membrane vesicles with %Lo≥50% [44]. The emission fluorescence in MFGM can reflect the lateral distribution of Lo, and the weaker light intensity implies the liquid-disordered (Ld) phase domain. Therefore, 3DOM can speculate the heterogeneity in the lateral organization of polar lipids.

### 3DOM imaging of SYTOX Orange-labeled λ-DNA

The binding of small dye molecules potentially induces structural changes in DNA [45], which can lead to altered DNA stability [46] and affect DNA processing of proteins [47]. A detailed understanding of the binding modes of fluorescent dyes to DNA is fundamental to drugs development, therapeutic and sensing applications. We prepare lambda phage DNA (*λ*-DNA) strands adhered to glass coverslips and used SYTOX Orange dye for labeling (see Methods section). A model of the labeling is shown in Fig. 6d. The samples are imaged at 65× 65 *μ*m^2^ fields of view at 16.67 fps.

**Fig. 6.**
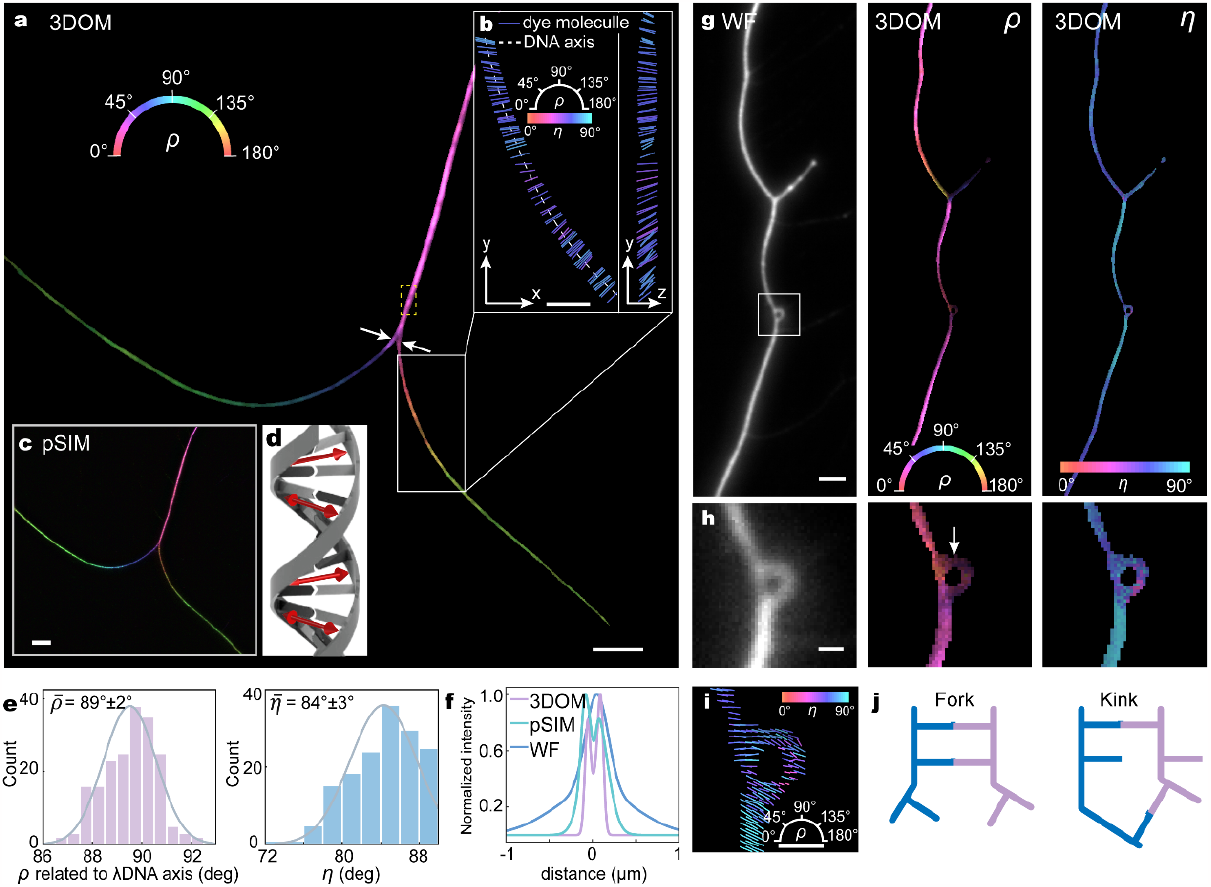
3DOM imaging of λ-DNA labeled with SYTOX Orange. **a** 3DOM images (*ρ* component) of λ-DNA strands with color coding based on the average azimuthal orientation of the dye molecules measured within 65 nm voxels. **b** Visualization of orientation measurements along a short strand of λ-DNA in *x-y* and *y-z* views (gray box in **(a)**). The rods point in the direction of *ρ* and color-coded to denote *η*. **c** pSIM images of λ-DNA, consistent with 3DOM results. **d** SYTOX Orange has an intercalation binding mode in DNA with an absorption dipole moment perpendicular to the DNA axis. **e** Histograms of orientation measurements along the DNA axis (yellow dash box in **(a)**). **f** Comparing 3DOM, pSIM and Widefield resolutions (white arrows in **(a)**). **g** Super-resolution images of λ-DNA strands exhibiting “bends”. **h** Zoom in on the *x-y* view of the boxed regions in **(g). i** Kinks formation occurs in DNA axis bending and the orientation of the visualized dye molecules changes (see arrows). **j** Possible state models for the localized structure of bending DNA. Scale bars (**a, c**), 5 μm; (**b, g**), 2 μm; (**h, i**), 500 nm.

Fig. 6a illustrates 3DOM super-resolution imaging of *λ*-DNA with a resolution of 128 nm (Fig. 6f), and our azimuthal results are consistent with those obtained by the pSIM method (Fig. 6c). We observe that the overall dye molecules are aligned perpendicular to the DNA strands, and as the DNA axis bends, the average orientation of the dye molecules rotates accordingly (Supplementary Video S2). Furthermore, in the *y-z* view we find that the angle *η* between the dye molecules and the *z*-axis is variable, showing a distribution similar to the helical shape of the DNA (Fig. 6b). By fitting the extension of the DNA axis, we histogram the orientation of the dye molecules relative to the DNA axis, and the *z*-axis, respectively (Fig. 6e). According to the histogram statistics, the fluorescent molecules are mostly perpendicular to the DNA axis (Δ*ρ* = 89°± 2°), and the precisions of *ρ* and *η* are obtained as 2° and 3°, respectively. The experimental precision is slightly greater than the precision of our measurement method, which may be due to the presence of secondary binding modes (secondary/major grooves, or DNA phosphate backbone) [9]. Our orientation measurements confirm that the STOYOX Orange fluorophores are inserted between neighboring bases of DNA such that the absorption dipole moment is aligned perpendicular to the DNA axis.

Since the SYTOX Orange dye molecules are oriented perpendicular to the DNA axis, they can be used to detect induced deformations within the DNA strand, where DNA bending involves many processes of gene expression [48]. Fig. 6g, h shows the bending of the DNA as it is deposited on the coverslip. It has been observed that even bare DNA can spontaneously bend into a circle [49]. By visualizing regions of high curvature (Fig. 6i), we observe that the orientation of the molecules presents a “non-perpendicular” situation. In this case, we believe that kinks are formed in DNA bending to release the bending stress stored in the DNA strands, and thus incomplete base pairing or incomplete stacking occurs. Referring to existing models of DNA bending [50-52], we speculate on two modes of kinking in this DNA bending (Fig. 6j). The difference between the two models is the state of base stretching [52]. Due to localized melting in the central region, base stretching occurs, and when “breathing” occurs at the end of the double strand, the fork state is formed.

### 3DOM imaging of actin filaments in U2OS cells

A comprehensive understanding of the intracellular distribution of actin filament orientations, along with the significant presence of 3D orientations, is crucial for explaining cellular motility powered by the actin cytoskeletal system [53]. We prepare samples of actin filaments immobilized on glass coverslips and labeled with Alexa Fluor 488 (AF488)-phalloidin (see Methods section), which acts as dipole attached to the target proteins (Fig. 7b).

**Fig. 7.**
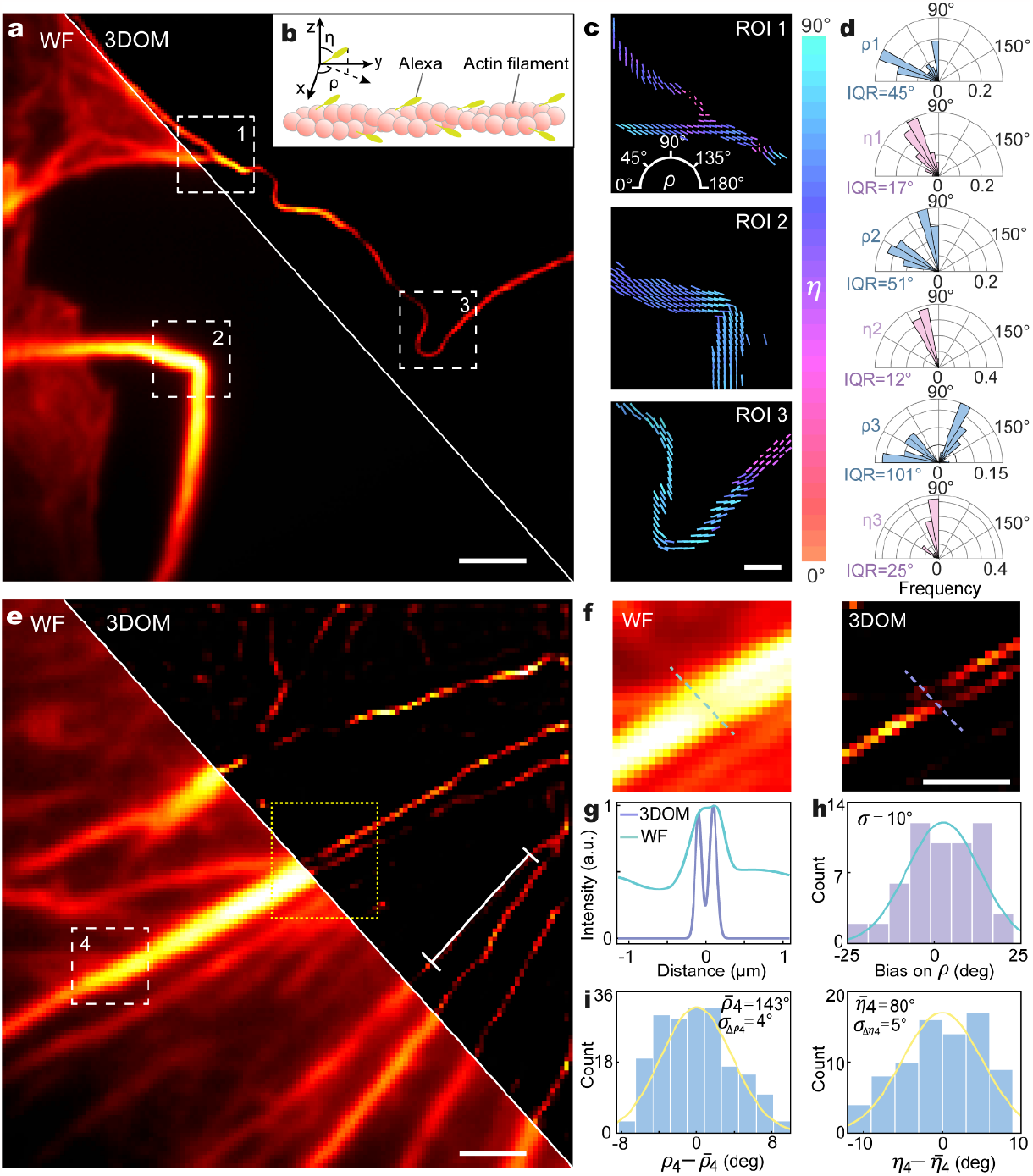
3DOM imaging of actin filaments in U2OS cells. **a** Comparison of WF and 3DOM imaging of actin filaments in U2OS cells. **b** The schematic diagram of the structure of actin filaments labeled by AF488-phalloidin, where the ensemble dipole orientation is parallel to the filament. The orientation of the AF488-phalloidin is defined by (*ρ, η*) in the coordinate system. **c** 3DOM results within three white dashed boxed in (**a**). The rods represent the orientation and are color-coded according to the measured *ρ* and *η*, respectively, with their lengths are proportional to sin *η*. **d** The distribution of azimuthal angle *ρ* and polar angle *η* in the region indicated in (**c**). **e** WF imaging and 3DOM imaging of the near-linear distribution of actin filaments. **f** The intensity distribution comparison between WF and 3DOM within the yellow dashed box in (**e**). **g** 3DOM can distinguish two filaments that cannot be resolved by WF imaging in (**f**). **h** The dipole histogram of the white line in (**e**), the deviation on *ρ* represents the difference between the dipole azimuth orientation and the filament direction. **i** Mean azimuthal value 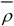 and polar value 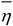 in the white dashed rectangular region shown in (**e**), along with the corresponding standard deviations *σ*_Δ*ρ*_ and *σ*_Δ*η*_, where 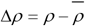 and 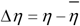. Scale bars (**a, e**), 1 μm; (**c**), 200 nm.

Figs. 7a, e show the distribution of actin filaments that potentially exhibit various morphologies as a result of localized mechanical stresses, such as bending, stretching, and torsion [54]. The intensity profiles in Fig. 7f, g demonstrate that 3DOM can resolve two filaments with a distance of 190 nm, which is not achievable in WF imaging. We use polar histograms to count the orientation distributions (Fig. 7c, d), and at the non-crossing of the cell edges, the filaments are adherent to the coverslip, thus *η* of ∼90° in ROI2 and ROI3, whereas there is a certain height of inclination of the filaments closer to the cells. The quantification in Fig. 7h shows an orientation deviation of 10° between the AF488-phalloidin dipoles and the actin filaments, demonstrating the azimuthal precision of 3DOM in measuring the molecular structure of actin filaments.

The angular distribution range of fluorophore labeling is useful for estimating local structural disorder of actin filaments [55]. Here, we use 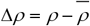 and 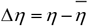 to quantify the local structural disorder, where 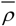 and 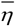 are the averaged value over the concerned region, is considered as the direction of the actin filament. The standard deviation *σ*_Δ*ρ*_ and *σ*_Δ*η*_ represent the spread of the probe orientations relative to the actin filament direction. By employing the resolved 3D orientation of the molecules, we compare the structural disorder of the filaments in the three regions (Supplementary Fig. S4 and Supplementary Table S2 in Supplementary Section 5), which is challenging to discern differences in these structural disorders only in the WF intensity diagram. In Fig. 7i, we obtain the average orientation distribution of actin filaments with Δ*ρ*_4_ and Δ*η*_4_ ranges of ∼20°, and standard deviations for Δ*ρ*_4_ and Δ*η*_4_ of ∼4° and ∼5°, respectively. The obtained local structural disorder Δ*ρ* is comparable to the average range of orientations derived in previous studies [55]. This finding further supports that actin filaments can form linear structures with collective orientation, which presumably relates to mechanical stress in the cell.

### 3DOM imaging of GFP-labeled microtubules in live U2OS cells

To demonstrate the live cell imaging capability of 3DOM, we perform dynamic imaging of microtubules in live U2OS cells expressing green fluorescent protein (GFP) (Fig. 8 and Supplementary Video S3). Here we capture microtubule images at 3.6 fps for a field of view of 27 × 27 *μ*m^2^ (Fig. 8a). The imaging speed is faster than the movement of the sample, avoiding motion-induced image blurring. We statistically count the *ρ* and *η* of the GFP combinatorial dipole for the two morphologies of microtubules (Fig. 8d, e). The projected orientation of the GFP ensemble dipoles in the *x-y* plane are mostly perpendicular to the microtubule filaments (Fig. 8d), which is consistent with previous reports [27]. Furthermore, we find that the GFP ensemble dipoles are not parallel or perpendicular to the *x-y* plane, which breaks our perception that microtubules are centrally symmetric (Fig. 8f).

**Fig. 8.**
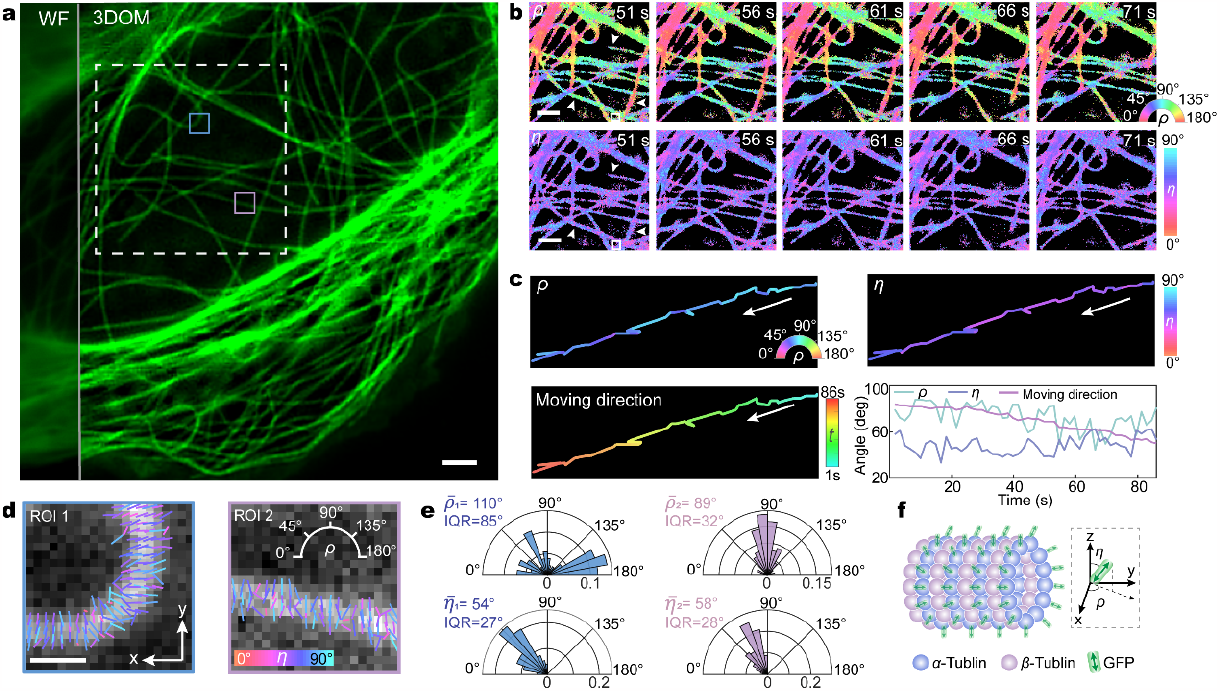
3DOM dynamic imaging of microtubules in live U2OS cells. **a** Comparison of WF and 3DOM imaging of live U2OS cells expressing tubulin-GFP. **b** Representative time-lapse image of the white dashed box region in (a), with the change in *ρ* shown in the upper part and the change in *η* shown in the lower part. **c** Time-lapse orientation and position of the microtubules in the white box in **(b). d** 3D orientation of GFP molecules (superimposed to WF images) within the pink and blue boxes in **(a)**, and the direction and color of the rods encode azimuthal angle *ρ* and polar angle *η*, respectively. **e** Polar histogram of the orientation of all GFP molecules in **(b). f** Schematic diagram of the tubulin-GFP structure. The projection direction of the ensemble dipole in the *x-y* plane is perpendicular to the microtubule filaments, and the ensemble dipole has an angle with the *z* axis. Scale bars (**a, b**), 2 μm; (**d**), 500 nm.

Fig. 8b shows a typical super-resolution time-lapse image of a microtubule over a period of 20 s. The growth and shortening process of microtubules can be directly seen at the white arrows. We further measure the orientation change of the microtubule endpoints during the growth and shortening process. In this process, the sliding direction is consistent with the measured azimuth angle *ρ*, and the change in polar angle *η* usually occurs when the microtubule undergoes a switch between shortening and growth (Fig. 8c). Studies show that microtubules and their associated motor proteins act in concert with actin and myosin to promote membrane tubulation [56]. Thus, 3DOM provides a new tool for observing microtubule stability and synergism, and offers insights into the dynamic behaviors of microtubules in vivo in an orientation dimension.

## Discussion

In this study, we have demonstrated 3DOM, for 3D orientation mapping of fluorescent molecules in WF that breaks the temporal-spatial resolution limitations. We demonstrate that 3DOM can effectively distinguish a minimum lateral spatial distance of 50 nm by mapping azimuthal and polar angles in theoretical analysis. When imaging the binding of KK114 dye to MFGs, we reconstruct a hemispherical MFG with a radius of 4.25 *μ*m and observe that fluorescent molecules are parallel to the membrane surface with an angle greater than 45° to the *z*-axis. This observation raises the interesting possibility that KK114 could be used to study the heterogeneity of biological cell membranes and the interactions of molecules on membranes. Furthermore, we measure the binding patterns of ***λ***-DNA to the dye molecules in 3D with 128 nm resolution and 2° precision, and resolve the structure of DNA strands. It suggests that intercalating dyes can be used as exquisite sensors to study DNA-binding proteins at the molecular level. By analyzing differences in the distribution of dipole orientations during imaging of filamentary biological samples, we also identify molecular-scale features of structural disorder, which is crucial for quantitative investigations of spatial arrangement and local disorder. Additionally, our 3DOM imaging of microtubules in living cells provides a molecular dimension to the dynamic behavior of microtubules in vivo. This knowledge serves as a reference point for unraveling the organization of filaments in future studies involving more complex and unknown assemblies.

Compared with existing FPMs, 3DOM has a high spatiotemporal resolution in resolving the 3D dipole orientation (see Supplementary Table S3 in Supplementary Section 6 for specific comparisons). Benefitted from the digital modulation of the SLM, our experimental setup allows for a raw image acquisition speed of 1697 fps and a reconstruction speed of 283 fps, ensuring efficient data acquisition and processing. While our current 3DOM methods employ a relatively simple model to improve computational speed, we acknowledge that for more complex samples, additional parameter descriptions need to be included in existing models to ensure accurate resolution [57, 58]. Considering the flexibility and mobility of the probe molecule in the biological structures, we complement the dipole tensor angle for subsequent updates of the 3DOM model (see Supplementary Section 7 for details). We also quantify the lower limits of precision using theoretical simulations of the CRB, where near-isotropic precision allows for reliable analysis of dipoles with unknown orientation. Additionally, we consider the critical case of oblique incidence. Exceeding the critical angle, the incident light undergoes a phase change, and it changes from linear polarization to elliptical polarization. To ensure incident light polarization, we recommend an oblique angle range of 15° to 60° based on simulation results. Taking into account the effect of equipment on polarization, we also emphasize that the extinction ratio must be higher than 39:1.

In summary, our proposed 3DOM method effectively overcomes the limitations of FPM in special resolution and 3D orientation mapping using WF imaging. 3DOM provides a more comprehensive understanding of the 3D spatial structure of fluorophore molecules. This enables us not only to distinguish various cytoskeletal organizations (actin filament and microtubule) but also to gain valuable insights into filament binding compactness, meshwork structures, and the extent of filament looseness. Moreover, 3DOM holds significant potential in DNA bending and the orientation of membranous organelles. One of the key advantages of 3DOM is its ease of upgradability to existing WF systems. The simple implementation, accurate 3D dipole orientation information, and superior spatiotemporal resolution of 3DOM makes it suitable for a wide range of applications, enhancing its accessibility and usability in different research settings. This powerful tool empowers researchers to unravel the intricate complexities of subcellular structure, biomechanics, and biodynamics, revolutionizing our understanding of cellular processes. We foresee 3DOM advancing understanding across a multitude of biological structures and interactions operative at the nanoscale.

## Methods

### Experimental setup of 3DOM

The optical system (Fig. 1a), involves three linearly polarized continuous-wave lasers,488 nm (Sapphire LP 488-200 mW, Coherent), 561nm (MGL-FN-561-200 mW, CNI) and 638 nm (WCP638-150FS-2108, PIC) as the excitation light source. The laser beam is coupled through a single-mode polarization maintaining fiber. Collimation of the beam is achieved using a lens1 (L1, AC254-150-A-ML, Thorlabs), after which it is directed towards a half-wave plate (HWP1, WPZ2420-400-650, Union Optic) and a polarizing beam splitter (PBS, CCM1-WPBS254/M, Thorlabs). Next, the outgoing light is projected onto a spatial light modulator (SLM, QXGA-R11, ForthDD), which modulates the light to generate a grating diffraction effect. The reflected light from the SLM passes through HWP2 and PBS once again before being focused by lens2 (L2, AC254-400-A-ML, Thorlabs). At the focal plane, a customized HWP and a mask are positioned. The customized HWP further modulates the outgoing light into p-polarized light, and the mask selectively allows only the +1st-order light to pass through. The transmitted light then proceeds through a relay lens comprising lens3 (L3, AC254-200-A-ML, Thorlabs) and lens4 (L4, AC254-300-A-ML, Thorlabs). These lenses converge the light at the back focal plane of the objective lens of the microscope (Eclipse Ti2, Nikon). The emitted fluorescence from the sample is collected through a polarization compensating dichroic mirror (ZT405-415/488/561/640rpc-phase R-UF3, Chroma). Finally, the fluorescence is captured by a camera (ORCA-Flash4.0 V3, Hamamatsu Photonics) for further analysis and imaging. This experimental setup enables precise control and manipulation of the light path, facilitating efficient data acquisition and analysis in the 3DOM method.

To account for non-uniform illumination at different angles during the experiment, we implement intensity calibration using a fluorescent slide as a sample. A set of intensity results is collected from the calibration sample, and the ratio between this calibration set and its mean is used as the correction factor to achieve intensity calibration for the sample results at corresponding angles. This ensures accurate and reliable dipole orientation solutions, improving the overall precision of our 3DOM method.

### Sample fabrication and preparation

#### Milk fat globule

We used fresh milk collected from farms and stored at -20 °C. A dilution mixture was created following the milk: ddH2O ratio of 1:20. Subsequently, we combined KK114 (STRED-0199-1MG, Abberior) with ddH2O in a ratio of 1:16. Then, the diluted milk and dye solution were mixed in a 1:1 ratio and reacted at 25°C for 15 min. Next, low-melting-point agarose was mixed with water in a 1:100 mass-to-volume ratio and heated to 75°C until the agarose granules were completely melted. Two clean glass slides were stacked on top of each other maintaining a thickness of ∼1 mm, then 33 μL of 1% agarose gel at 75°C was dropped between the slides and left to solidify for 30 min, and gently separated using forceps. Finally, we placed 5 μL of the liquid sample into an imaging dish, covered it with a gel pad, and fixed it with nail polish. Note that all procedures should be carried out at room temperature (25°C).

#### λ-DNA Sample

Poly Methyl Methacrylate (0.5g, M9810, Solarbio) was dissolved in 10 mL denture water. After that, the supernatant was extracted and applied to a slide to form a film. We diluted 0.3 μL λ-DNA (SD0021, Invitrogen) and 32 μL 1000×SYTOX™ Orange Nucleic Acid Stains (Invitrogen, S11368) in 968 μL PBS. Then, we applied 1 μL of the solution to the slide and let it dry. The coverslip was sealed on the slide with the ProLong Diamond mountant (P36965, Invitrogen).

#### Labeling of actin in U2OS cells

Upon reaching 75% confluence, the U2OS cells were fixed for 15 min at room temperature using a 4% formaldehyde solution (Invitrogen, R37814). Subsequently, the cells were permeabilized with 0.1% Triton™ X-100 (HFH10, Invitrogen) in PBS for 5 min, followed by rinsing with PBS to remove residual Triton™ X-100. To label the actin filaments, the cells were stained with AF488-Phalloidin (A12379, Invitrogen) dye for 1 h at room temperature. After staining, the samples were washed twice with PBS to remove any excess dye. The coverslips carrying the stained cells were then air-dried in the dark. Finally, 30 μL of ProLong (P36984, Invitrogen) mounting medium was added to each coverslip, and left to air-dry overnight at 4°C.

#### Plasmid Transfection

For GFP labeling of tubulin, the pc3.1-GFP-tubulin plasmid was transfected into U2OS cells under the standard protocol of Lipofectamine 3000 (L3000, Invitrogen). Briefly, transfection experiments were performed until the density of U2OS cells reached around 80%, and 1 μg plasmid was transfected into the cells. After transfection for 6 hours, replaced with fresh cell culture medium, and cultured for around 24 hours before performing super-resolution imaging.

#### Cell Maintenance and Preparation

U2OS cells were cultured Dulbecco’s modified Eagle’s medium (DMEM, 995-040, Gibco), which was a high-glucose medium supplemented with 10% bovine serum (10099, Gibco) and 1% penicillin-streptomycin antibiotic (10,000 U/mL, 15140148, Gibco). The cells were maintained in a controlled environment at 37 °C with a 5% CO_2_ atmosphere until they reached ∼75% confluency. For fixed-cell imaging experiments, the cells were seeded onto coverslips (CG15CH2, Thorlabs).

## Code availability

The codes for experimental data acquisition, physical model simulation, and Cramér-Rao bound calculation are available from the corresponding author.

## Data availability

The associated datasets are available from the corresponding author upon reasonable request.

## Acknowledgments

We thank Dr. Chongqi Zhou for providing Cramér-Rao bound theoretical support. This work is supported by the National Key R&D Program of China (2022YFC3401100) and the National Natural Science Foundation of China (62335008, 62025501, 31971376, 92150301). We thank Airy Technology Co. Ltd., for providing with Airy Polar-SIM system to build up 3DOM system and image.

## Author contributions

K.Z., M.L. and P.X. conceived the project. S.Z. performed the experiments and the simulations with the help of L.Q. and X.G.. X.X. and K.Z. assisted with the theoretical modelling. Y.F., S.G. and H.H. prepared the samples. M.L. helped to supervise the experiments and interpret the results. W.W. assisted with the analysis of the results. S. Z. and P.X. wrote the manuscript with contributions from all authors.

